# Identification of a chromatin-bound ERRα interactome network in mouse liver

**DOI:** 10.1101/2024.01.02.573907

**Authors:** Charlotte Scholtes, Catherine Rosa Dufour, Emma Pleynet, Samaneh Kamyabiazar, Phillipe Hutton, Reeba Baby, Christina Guluzian, Vincent Giguère

## Abstract

**Objective:** Estrogen-related-receptor α (ERRα) plays a critical role in the transcriptional regulation of cellular bioenergetics and metabolism, and perturbations in its activity have been associated with metabolic diseases. While several coactivators and corepressors of ERRα have been identified to date, a knowledge gap remains in understanding the extent to which ERRα cooperates with coregulators in the control of gene expression. Herein, we mapped the primary chromatin-bound ERRα interactome in mouse liver.

**Methods:** RIME (Rapid Immuno-precipitation Mass spectrometry of Endogenous proteins) analysis using mouse liver samples from two circadian time points was used to catalog ERRα-interacting proteins on chromatin. The genomic crosstalk between ERRα and its identified cofactors in the transcriptional control of precise gene programs was explored through cross-examination of genome-wide binding profiles from chromatin immunoprecipitation-sequencing (ChIP-seq) studies. The dynamic interplay between ERRα and its newly uncovered cofactor Host cell factor C1 (HCFC1) was further investigated by loss-of-function studies in hepatocytes.

**Results:** Characterization of the hepatic ERRα chromatin interactome led to the identification of 48 transcriptional interactors of which 42 were previously unknown including HCFC1. Interrogation of available ChIP-seq binding profiles highlighted oxidative phosphorylation (OXPHOS) under the control of a complex regulatory network between ERRα and multiple cofactors. While ERRα and HCFC1 were found to bind to a large set of common genes, only a small fraction showed their co-localization, found predominately near the transcriptional start sites of genes particularly enriched for components of the mitochondrial respiratory chain. Knockdown studies demonstrated inverse regulatory actions of ERRα and HCFC1 on OXPHOS gene expression ultimately dictating the impact of their loss-of-function on mitochondrial respiration.

**Conclusions:** Our work unveils a repertoire of previously unknown transcriptional partners of ERRα comprised of chromatin modifiers and transcription factors thus advancing our knowledge of how ERRα regulates metabolic transcriptional programs.

## 1. Introduction

Estrogen-related receptor α (ERRα) is an orphan member of the nuclear receptor superfamily of transcription factors that acts as a master regulator of gene programs involved in cellular energy metabolism, insulin sensitivity, and the stress response [1–3]. Key transcriptionally regulated metabolic gene programs include fatty acid metabolism, oxidative phosphorylation (OXPHOS), glycolysis, and generation of reactive oxidative species [4–12]. Accordingly, correlations have been observed between perturbations in the levels of genes controlled by ERRα and diseases such as obesity, non-alcoholic fatty liver disease (NAFLD), diabetes, and certain types of cancer [3; 8; 13-15]. In a manner similar to other nuclear receptors, ERRα action as an activator or repressor of gene expression requires interactions with a specific set of coregulator proteins [16]. The coregulators peroxisome proliferator-activated receptor γ co-activator 1 α and β (PGC-1α/β) have been shown to strongly enhance the transcriptional activity of ERRα [17–20] while prospero homeobox protein 1 (PROX1) represses its activity [9]. Other coregulators such as the receptor-interacting protein 140 (RIP140) can function as either coactivator or corepressor for ERRα depending on the genomic targets [21]. Thus, the identification of ERRα chromatin-bound partners is important for our understanding of the molecular mechanisms underlying energy cellular metabolism and their potential implication in the development of of metabolic disorders.

In this study, the Rapid Immunoprecipitation Mass spectrometry of Endogenous proteins (RIME) technique [22; 23] was employed to elucidate the ERRα chromatin-bound interactome. By combining chromatin immunoprecipitation (ChIP) with mass spectrometry, RIME enables the identification of proteins that can either form a complex with the bait protein through direct or indirect interactions as well as those that bind in proximity on chromatin. Herein, we report the identification of a network of 48 ERRα-interacting proteins, of which 42 were previously unknown. Further interrogation of ChIP-sequencing (ChIP-seq) profiles uncovered OXPHOS-related genes at the hub of a gene regulatory network between ERRα and of its cofactors, including the novel interactor HCFC1 (Host cell factor C1). Notably, the specific set of loci found shared by ERRα and HCFC1 was predominantly enriched for OXPHOS gene promoters. Genetic knockdown and functional studies in hepatocytes showed a dependency on HCFC1 for ERRα transcriptional control of mitochondrial respiration, highlighting the broader implications of ERRα’s interactome on hepatocyte physiology. Together, this work identified a comprehensive ERRα chromatin interaction network that sheds new insight on the molecular mechanisms underlying ERRα-mediated gene regulation and metabolic control that may prove beneficial for the development of therapeutic strategies for the treatment of metabolic disorders.

## 2. Materials and methods

### 2.1 Animals

All mouse experiments were conducted in accordance with established standards for animal care and all protocols received approval from the McGill Facility Animal Care Committee and the Canadian Council on Animal Care. Wild-type (WT, Envigo) in a C57BL/6NHsd genetic background were group-housed with two to five mice per cage. Mice were kept in an animal facility at McGill University and had ad libitum access to a standard rodent diet (2920X, Envigo, irradiated) and water. The animals were maintained under constant environment (ambient temperature 22°-24°C; relative humidity: 30%-70%) with a 12-hour light-dark cycle (7:00 a.m. to 7:00 p.m.). Male mice aged 2 months were used for all experiments. Mice were sacrificed by cervical dislocation and freshly isolated liver tissues were immediately processed for RIME or ChIP-qPCR studies.

### 2.2 Cell culture

Hepa 1-6 cells, originally sourced from the ATCC (RRID: CVCL_0327), were maintained in DMEM (319-005-CL, Wisent) supplemented with 10% (v/v) fetal bovine serum (FBS, 12483020, Thermo Fisher Scientific) and 100 units/ml penicillin-streptomycin (450-201-EL, Wisent). Cultured cells were maintained at 37°C with 5% CO_2_, and mycoplasma contamination was regularly assessed using the polymerase chain reaction (PCR) mycoplasma detection kit (G238, ABM).

### 2.3 Nuclear extraction for RIME

Mice (n=3) were euthanized at 2 months of age by cervical dislocation at both Zeitgeber Time (ZT) 8 and ZT24 corresponding to the time of lights on and off, respectively. The livers were quickly isolated and homogenized in 10 mL of cold PBS in 15 ml conical tubes, followed by centrifugation at 3,000 rpm for 2 min at 4°C. The resulting cell pellets were then incubated with room temperature PBS containing 2 mM DSG (disuccinimidyl glutarate, sc-285455, Santa Cruz Biotechnology) for 25 min with gentle agitation. The 2 mM DSG/PBS solution was freshly prepared from a 1 M DSG stock solution in DMSO stored at −20°C. Following protein-protein crosslinking with DSG, formaldehyde was added to a 1% v/v final concentration (2106-01, JT Baker) to crosslink protein-DNA and the samples were left for 20 min at room temperature with mild agitation. Crosslinking was then quenched by the addition of glycine (0.1 M, pH 3.5). The cell pellets were washed twice with ice-cold PBS with centrifugation at 3,000 rpm for 2 min at 4°C after each rinse. Following the rinsing steps, 6 ml of cold lysis buffer 1 (LB1: 50 mM Hepes-KOH pH 7.5, 140 mM NaCl, 1 mM EDTA, 10% (v/v) glycerol, 0.5% NP-40/Igepal CA-630, 0.25% Triton-X-100) was added, and the samples were left to rotate at 4°C for 10 min. The lysates were then cleared by centrifugation at 2,000g for 5 min at 4°C. Subsequently, the pelleted nuclei were resuspended with 10 ml of cold LB2 (10 mM Tris-HCl pH 8.0, 200 mM NaCl, 1 mM EDTA, 0.5 mM EGTA), and the samples were left to rotate at 4°C for 5 min. The samples were again cleared by centrifugation at 2,000g for 5 min at 4°C. Next, the purified nuclear pellet from each liver was resuspended in 2 ml of LB3 (10 mM Tris-HCl pH 8.0, 100 mM NaCl, 1 mM EDTA, 0.5 mM EGTA, 0.1% (wt/v) sodium deoxycholate, 0.5% (v/v) N-lauroylsarcosine). Each nuclear sample was aliquoted into 500 µl portions in Eppendorf tubes for subsequent sonication (Fisher Scientific, model 100; Power=11, 40 x 8 sec pulses) in an ice/water bath until the solution became clear. To enrich the sonicated nuclei, 30 µL of 10% Triton X-100 was added to each sample, and the samples were vortexed and then centrifuged at 13,000g for 15 min at 4°C. The resulting supernatants were transferred to new tubes. The efficiency of sonication was assessed by agarose gel electrophoresis. For this purpose, 20 µl of sonicated nuclear lysate was added to 200 µl of chromatin decrosslinking buffer (1% SDS, 0.1 M NaHCO3) with 8 µl of 5M NaCl and 10 µL of 20 mg/ml proteinase K. This mixture was incubated at 65°C for 1 h for rapid decrosslinking. The decrosslinked samples were then purified using a QIAquick PCR purification kit (28106, Qiagen), and the samples were eluted with 30 µl of homemade elution buffer (10 mM Tris-HCl pH 8.0, 0.1 mM EDTA pH 8.0). Of the 30 µl eluate, 10 µl was loaded onto a 1% agarose gel with ethidium bromide to verify the sonication efficiency. Chromatin was considered well sheared when DNA fragments were mostly between 100-1000 bp with an enrichment at 500 bp. DNA concentration was measured using a Nanodrop. It is important to note that all buffers used in the RIME experiments were supplemented with protease and phosphatase inhibitors (A32955, A32965, Thermo Fisher) before use and were stored at 4°C.

### 2.4 Chromatin immunoprecipitation (ChIP) for RIME

For each ChIP assay, 50 μl of Dynabeads® Protein A (10008D, Invitrogen) underwent four washes, each with 1 ml of ice-cold PBS containing 5 mg/ml of bovine serum albumin (BSA, 800-095-CG, Wisent). Subsequently, the beads were resuspended in 500 μl of ice-cold PBS/BSA, to which either 0.8 μg of anti-ERR antibody (ab76228, Abcam; RRID: AB_1523580) or 0.8 μg of non-specific rabbit IgG (10500C, Thermo Fisher; RRID: AB_2532981) was added. The antibody-bead mixtures were then placed on a rotator and incubated overnight at 4°C. The following day, the antibody-bound beads were washed four times with 1 ml of ice-cold PBS/BSA to eliminate any unbound antibody. Subsequently, 200 μg of sonicated nuclear lysate was added, and the tubes were left to rotate at 4°C overnight. On the subsequent day, the beads were washed three times with 1 ml of ice-cold RIPA buffer (50 mM Hepes pH 7.6, 0.5 M LiCl, 1 mM EDTA pH 8.0, 0.7% (wt/v) sodium deoxycholate, 1% (v/v) NP-40). The beads then underwent an additional four washes, each with 1 ml of a cold 100 mM ammonium bicarbonate (AMBIC, 14249, Alf Aesar) solution to remove detergents and salts. During the second AMBIC wash, the beads were transferred to a new tube. To complete the process, 50 μl of 100 mM AMBIC solution was added to the beads prior to trypsin digestion.

### 2.5 On-beads digestion and LC-MS/MS for RIME

On-beads trypsin digestion was performed overnight at 37°C with a ratio enzyme:protein of 1:20 in 2 M urea/50 mM ammonium bicarbonate. The samples were then reduced with 13 mM dithiothreitol at 37°C and, after cooling for 10 min, alkylated with 23 mM iodoacetamide at room temperature for 20 min in the dark. The supernatants were acidified with trifluoroacetic acid 100% and cleaned from residual detergents and reagents with MCX cartridges (Waters Oasis MCX 96-well Elution Plate) following the manufacturer’s instructions. Samples were dried with a Speed-vac and stored at −20°C for a few days. Samples were reconstituted under agitation for 15 min in 12 µl of 2% acetonitrile/1% formic acid and loaded into a 75 μm i.d. × 150 mm Self-Pack C18 column installed in the Easy-nLC II system (Proxeon Biosystems). The HPLC system was coupled to a LTQ Orbitrap Velos spectrometer (Thermo Scientific) through a Nanospray Flex Ion Source. The buffers used for chromatography were 0.2% formic acid (solvent A) and 90% acetonitrile/0.2% formic acid (solvent B). Peptides were eluted with a three-slope gradient at a flowrate of 250 l/min. Solvent B first increased from 2 to 29% in 70 min, then from 29 to 44% in 50 min and finally from 44 to 95% B in 6 min. LC-MS/MS data acquisition was accomplished using a seventeen-scan event cycle comprised of a full scan MS for scan event 1 acquired in the Orbitrap. The mass resolution for MS was set to 60,000 (at m/z 400) and used to trigger the sixteen additional MS/MS events acquired in parallel in the linear ion trap for the top sixteen most intense ions. Mass over charge ratio range was from 360 to 1,700 for MS scanning with a target value of 1,000,000 charges and from ∼1/3 of parent m/z ratio to 2000 for MS/MS scanning with a target value of 10,000 charges. The data-dependent scan events used a maximum ion fill time of 100 ms and 1 microscan. Target ions already selected for MS/MS were dynamically excluded for 31 s after 2 counts. Nanospray and S-lens voltages were set to 1.3–1.7 kV and 50 V, respectively. Capillary temperature was set to 250°C. MS/MS conditions were normalized collision energy, 35 V; activation q, 0.25; activation time, 10 ms.

### 2.6 RIME protein identification

Peak list files were generated using Proteome Discoverer (version 2.3) (OPTON-20141, Thermo Scientific) with the following parameters: a minimum mass of 500 Da, a maximum mass of 6000 Da, no grouping of MS/MS spectra, automatic setting for precursor charge, and a minimum of 5 fragment ions. Protein database searching was conducted with Mascot 2.6 (Matrix Science) against the mouse Uniprot complete protein database (dated April 15th, 2015). The mass tolerances for precursor and fragment ions were set at 10 ppm and 0.6 Da, respectively. Trypsin was designated as the enzyme, allowing for up to 2 missed cleavages. Cysteine carbamidomethylation was set as a fixed modification, and methionine oxidation as a variable modification. Data interpretation was carried out using Scaffold (version 4.10.0).

### 2.7 RIME data processing and functional analysis

Protein quantification based on peak ion intensity, as well as the integration and normalization of the samples, was conducted using the proteome software tool Scaffold v4.10.0. The thresholds 1% protein and 1% peptide false discovery rate (FDR) were applied with a minimum requirement of 2 detected peptides for inclusion. Normalization based on total precursor ion (TPI) intensity values was performed across all samples and subsequent computational analyses were conducted outside of Scaffold. Briefly, quantitative values were log2-transformed, missing values were treated as “NaN,” and protein clusters and DECOYs were excluded. For each experimental condition (ZT8 and ZT24), proteins were retained only if peptides were detected in at least 2 out of 3 replicates, and we could confirm the presence of 2 unique peptides in at least 1 replicate. Only proteins with no peptides found in the matching IgG controls or with a mean TPI fold-change ≥ 5 over IgG were retained. Quantitative values for the identified RIME ERRα chromatin-interacting proteins in mouse livers harvested at ZT8 and ZT24 and their comparison are summarized in Suppl. Table 1.

The identified list of 48 ERRα interactors by RIME was analyzed by Metascape v3.5.20230501 [24] and Ingenuity Pathway Analysis (IPA, Qiagen) v 01-22-01 Fall 2023 for enrichment analysis of biological terms and canonical pathways, respectively, using gene names as input with default settings. For IPA, Fisher’s exact test p-values associated with enriched canonical pathways were corrected for multiple testing (adjusted p-value, Padj) using the Benjamini-Hochberg method.

Protein-protein interaction networks for the ERRα interactors identified at ZT8 (28) or ZT24 (40) were constructed using IPA’s build tool with direct and indirect protein connections based on IPA’s knowledgebase.

The functional protein-protein interaction network for the total 48 ERRα interactors found was generated using Metascape with default settings. The Molecular Complex Detection (MCODE) algorithm [25] was applied to detect subcomplexes with highly interconnected factors and their functional attributes.

Normalized mouse and human expression data (transcripts per million, TPM) of normal liver and NAFLD were downloaded from GepLiver [26] to measure using Spearman correlation the strength and degree in correlation between *Esrra* mRNA levels and the expression of a subset of its identified RIME interactors. Mouse (normal, n=94, NAFLD, n=125) and human (normal, n=362; NADLD, n=503).

### 2.8 Compilation of previously known ERRα interactors

Known ERRα-interacting proteins were compiled from four independent sources: BioGRID v4.4.227 (https://thebiogrid.org) [27], HuRI (http://interactome.baderlab.org/) [28], IID (http://ophid.utoronto.ca/iid) [29], and STRING v12.0 (https://string-db.org) [30]. For BioGRID, both human and mouse ERRα interactors were considered from all high and low throughput assays. HuRI was limited to human protein-protein interactions and for IID, both human and mouse ERRα interactors were retrieved from experimentally determined interactions. For STRING, both human and mouse ERRα interactors were considered from experimentally determined interactions only using a minimum low confidence score of 0.150. A total of 112 ERRα interactors from these four sources are listed in Suppl. Table 2.

### 2.8 siRNA experiments

In the siRNA-mediated knockdown experiments targeting ERRα and HCFC1, Hepa 1-6 cells were initially cultured in DMEM medium. Subsequently, the cells were trypsinized, and 400,000 cells were reseeded in 6 cm plates. Cells were then transfected with 100 nM of either a control pool of scrambled siRNAs (D-001810-01-50, Dharmacon) or siRNAs targeting mouse ERRα (L-040772-00-0050, Dharmacon) or HCFC1 (L-051186-01-0050, Dharmacon). To elaborate, 4 μl of siRNAs were mixed with 37.5 μl of Hiperfect transfection reagent (301707, Qiagen) and enough Optimem Opti-MEM reduced serum medium (31985070, Thermo Fisher) to reach a final volume of 1 ml. This mixture was allowed to incubate at room temperature for 10 min then added to the cells, and media was added to have a final volume of 4 ml, bringing the final siRNA concentration to 100 nM. The siRNA transfection was repeated after 48 h, directly on the attached cells, and the cells were left for either another 24 h for determination of cell numbers or 48 h for RNA-seq, ChIP-qPCR, ChIP-seq, and Agilent Seahorse mitochondrial respiration experiments.

### 2.9 ChIP-qPCR

For ChIP-qPCR experiments involving mouse liver samples, one liver from a WT mouse was homogenized in cold 1X PBS containing protease inhibitors using a polytron. After centrifugation at 3000 rpm during 2 min at 4°C, the cell pellets were resuspended in 1X PBS containing 1% formaldehyde (AA33314-K2, VWR) and protease inhibitors, followed by rotation at room temperature for 12 min to facilitate DNA-protein crosslinking. Subsequently, the samples were centrifuged at 3000 rpm during 2 min at 4°C, and the cell pellets were washed twice with cold 1X PBS before being resuspended in 10 ml of cell lysis buffer (5 mM HEPES pH 8, 85 mM KCl, 0.5% NP-40, and protease inhibitors) and rotated at 4°C for 30 min with vortexing every 10 min. The nuclear pellets were collected after centrifugation at 3000 rpm for 10 min at 4°C. The nuclear pellets were then resuspended in 2 ml of nuclei lysis buffer (50 mM Tris-HCl pH 8.1, 10 mM EDTA, 1% SDS, and protease inhibitors), aliquoted in volumes of ∼700 μl in Eppendorf tubes and sonicated on ice until the DNA fragments were enriched to around 200–500 bp with a manual sonicator (model 100, Fisher Scientific). For ChIP assays, 10 μl of ERRα antibody (ab76228, Abcam; RRID: AB_1523580), 10 μl of HCFC1 antibody (A301-399A, Bethyl; RRID: AB_961012), or 1 μl of non-specific rabbit IgG (10500C, Thermo Fisher; RRID: AB_2532981) were pre-bound overnight at 4°C to 60 μl of Dynabeads protein G (10003D, Thermo Fisher Scientific) diluted in 250 μl of blocking buffer (PBS, 0.5% BSA, and protease inhibitors). The day after, 200 μg of chromatin DNA was diluted in 2.5X ChIP dilution buffer (2 mM EDTA pH 8.0, 100 mM NaCl, 20 mM Tris-HCl pH 8.0, 0.5% Triton X-100, and protease inhibitors) along with 100 μl of blocking buffer and then added to the antibody-bound beads. The beads were washed three times with blocking buffer and rotated overnight at 4°C. The beads were washed three times with 1 ml of cold LiCl wash buffer (100 mM Tris-HCl pH 7.5, 500 mM LiCl, 1% NP-40, 1% Na-deoxycholate), rotating at 4°C for 3 min per wash. Subsequently, the beads were transferred to a new tube and washed two more times with 1 ml of LiCl buffer, followed by a brief wash with 1 ml of cold TE buffer (10 mM Tris-HCl pH 7.5 and 1 mM EDTA pH 8.0). The beads were then subjected to decrosslinking in 300 μl of decrosslink buffer (1% SDS and 0.1 M NaHCO3) at 65°C overnight. The supernatant was collected and incubated with 300 μl of TE buffer and 0.2 μg/μl RNase A (19101, QIAGEN) at 37°C for 2 h, followed by incubation at 55°C with 0.2 μg/μl Proteinase K (PRK403.250, BioSHOP) for 2 h. The ChIP DNA was purified using a QIAquick PCR Purification Kit (28106, QIAGEN) and eluted with 40 μl of a homemade elution buffer (10 mM Tris-HCl pH 8.0 and 0.1 mM EDTA pH 8.0). ChIP-qPCR was performed on a LightCycler 480 instrument (Roche) using SYBR Green I Master Mix (4887352001, Roche). Specific primers for ChIP-qPCR analysis are listed in Suppl. Table 3. ChIP enrichment values were normalized against IgG controls and two negative control regions.

For ChIP-qPCR experiments in cells with siRNA-mediated knockdown, the Hepa 1-6 cell line was initially seeded in 6 cm cell culture dishes. Subsequently, cells were transfected with 100 nM of either a control pool of scrambled siRNAs (D-001810-01-50, Dharmacon) or siRNAs targeting mouse ERRα (L-040772-00-0050, Dharmacon) or HCFC1 (L-051186-01-0050, Dharmacon), as previously described above. The siRNA transfection was repeated after 48 h, directly on the attached cells. After a total of 96 h, cells from four plates for each condition were collected. For the ChIP procedure, the ChIP-IT High Sensitivity® kit (53040, Active Motif) was employed according to the manufacturer’s instructions with some modifications: 60 strokes were used for homogenization using a loose Dounce homogenizer and sonication was performed with 700 μl of ChIP buffer using an automatic sonicator (Qsonica Shearing System, model Q800R) set for 20 min of continuous operation with a 30-second ON/30-second OFF cycle at 40% amplitude. For each ChIP, 30 μg of chromatin was utilized, along with either 10 μl of HCFC1 antibody (A301-399A, Bethyl; RRID: AB_961012) or 1 μl of non-specific rabbit IgG (10500C, Thermo Fisher; RRID: AB_2532981). At the end of the ChIP procedure, the ChIP DNA was eluted in 100 μl of elution buffer. ChIP-qPCR was performed on a LightCycler 480 instrument (Roche) using SYBR Green I Master Mix (4887352001, Roche). Specific primers for ChIP-qPCR analysis are listed in Suppl. Table 3. ChIP enrichment values were normalized against IgG controls and two negative control regions.

### 2.10 ChIP-seq and data analysis

To validate and explore the chromatin associations between ERRα and its candidate partners identified by RIME analysis, we exploited the functionality of the Enrichment Analysis tool within ChIP-Atlas [31] that can establish the enrichment of transcriptional regulators at a given set of genomic coordinates using housed uniformly processed public ChIP-seq binding profiles. The analysis focused on the overlap of mouse (mm10) liver ERRα peak regions (SRX218535) with transcription factor ChIP-seq profiles available from any given tissue/cellular context based on peaks called using a q-value < 1E-05 with the default permutation (1x). The results were focused in on intersections between ERRα and its identified interactors from RIME, of which 24 of 48 interactors had available data. Among these refined results, 9 interactors of ERRα had available ChIP-seq profiles performed in liver tissue or hepatocytes. A summary of the results from the ChIP-Atlas Enrichment Analysis and selected liver/hepatocyte-specific datasets for further is shown in Suppl. Table 4. Liver/hepatocyte-specific ChIP-seq peak lists (q < 1E-05, mm10) downloaded from ChIP-Atlas were used for HOMER v4.11.1 [32] peak intersections using the script *mergePeaks* with default parameters (dgiven) and overlapping peaks were annotated using *annotatePeaks.pl*. Prior to ChIP-seq peak intersections, data from replicate experiments were compiled. ChIP-seq tracks were visualized using the UCSC Genome Browser (https://genome.ucsc.edu/). For KEGG pathway enrichment analysis (Human 2021) using EnrichR [33], genes with overlapping peaks found within ± 20kb of gene TSSs were used.

For an in-depth analysis of public ERRα and HCFC1 ChIP-seq data presented in Figure 5, the data were re-analyzed from raw fastq sequencing files: mouse liver ERRα (GSM1067408) [34], mouse liver HCFC1 (GSM3189034) [35]. Reads were mapped to the mouse reference genome (mm10) using BWA-MEM v0.7.12 [36] in paired-end mode at default parameters. Only reads that had a unique alignment (mapping quality > 20) were retained and PCR duplicates were removed using Picard tools v2.0.1 (https://broadinstitute.github.io/picard/). Peaks (summit +/-150 bp) were called using MACS2 software suite v2.1.1.20160309 [37] with the default FDR (alpha = 0.05) using sequenced libraries of IgG or input DNA as control for the ERRα and HCFC1 ChIP-seqs, respectively. Peaks in mitochondrial chromosome and scaffold regions were removed. HOMER peak intersections and annotations were performed as described above and following filtering of peaks ± 20kb of gene TSSs. The resulting set of 601 genomic regions annotated to 597 genes with ERRα and HCFC1 co-localization are shown in Suppl. Table 5. Separate “reference peak sets” was generated by merging ChIP-seq peaks across samples in the same experiment, using bedtools merge v2.27.0 with parameters:-sorted -d −150 (https://bedtools.readthedocs.io/). Peak signals were then calculated as Fragments Per Kilobase of transcript per Million mapped reads (FPKM) using HOMER. ChIP-seq tracks were visualized using the UCSC Genome Browser (https://genome.ucsc.edu/). Bigwig coverage files were created with the HOMER *makeUCSCfile* command and *bedGraphToBigWig* v4 utility from UCSC. Data were normalized so that each value represents the read count per base pair per 10^6^ reads. Heatmaps and average profiles were generated using modules “computeMatrix” (--referencePoint center) followed by “plotHeatmap” and “plotProfile” from deepTools v3.5.0 REF using bigwig files as input. De novo motif analysis was performed using HOMER with the *findMotifsGenome.pl* script within ± 150 bp of peak centers.

For HCFC1 ChIP-seq experiments in Hepa 1-6 cells, cells were treated with siRNAs against ERRα, HCFC1, or control siRNAs as described above. ChIPs were prepared as described in detail above for Hepa 1-6 cells and two independent ChIPs were pooled for ChIP-seq experiments. ChIP’d DNA and IgG controls for peak calling were prepared and sequenced using the NovaSeq 6000 platform (Illumina) as 150bp paired-end reads by Novogene Bioinformatics Technology Co., Ltd. Raw sequencing ChIP-seq data was processed as described above for re-analysis of public mouse liver ERRα and HCFC1 ChIP-seq datasets. HCFC1 ChIP-seq results from Hepa 1-6 cells ±ERRα or HCFC1 siRNA-mediated knockdown are presented in Suppl. Table 6.

### 2.11 RNA extraction and real-time quantitative PCR (RT-qPCR)

Total RNA was extracted from frozen Hepa 1-6 cell pellets using the RNeasy Mini kit (74106, QIAGEN) following the manufacturer’s instructions with an on-column DNase digestion step (79254, Qiagen). cDNA was made from 1 μg of RNA by reverse transcription with random primers (48190011, Fisher Scientific), dNTPs (10297-018, Thermo Fisher Scientific), RNase inhibitor (10777019, Fisher Scientific), 5X buffer, DTT, and ProtoScript® II reverse transcriptase (M0368L, New England Biolabs). RT-qPCR was performed on a LightCycler 480 instrument (Roche) using SYBR Green I Master Mix (4887352001, Roche). The relative expression of genes was normalized to the expression of *Tuba1a.* Specific RT-qPCR primer sequences are listed in Suppl. Table 3.

### 2.12 RNA-seq and data analysis

Total RNA was extracted as described above and used for Illumina mRNA sequencing by Novogene Bioinformatics Technology Co., Ltd. In brief, the RNA underwent quantification and quality assessment before library preparation using the NEBNext Ultra RNA Library Prep Kit for Illumina (NEB) in accordance with the manufacturer’s recommendations. The prepared libraries were sequenced on an Illumina platform (NovaSeq 6000) as 150 bp paired end reads. Hisat2 (version 2.0.5) was employed to align the paired-end clean reads to the mouse reference genome mm10. FeatureCounts (v1.5.0-p3) was used to count the number of reads per gene during the mapping process. The Fragments per Kilobase of Transcript Sequence per Million Base Pairs Sequenced (FPKM) for each gene were computed based on the gene’s length and the read counts mapped to it. Differential expression analysis between two groups (n=3 per group) was carried out using the DESeq2 R package (version 1.20.0) and genes with an adjusted p-value < 0.05 and log2|fold change| ≥ 0.5 were considered significantly altered. RNA-seq data is presented in Suppl. Table 7.

Gene set enrichment analysis (GSEA) was performed using GSEA software version 4.3.2 (https://www.gsea-msigdb.org/gsea/index.jsp) to identify enriched hallmark gene signatures within the Molecular Signature Database (MSigDB, mh.all.v2023.1.Mm) using Ensembl gene ID’s as input. Gene sets ranging from 15 to 2,000 genes were considered and all other parameters were based on default settings.

### 2.13 Protein extraction and Western blotting

For whole cell extracts from cultured Hepa 1-6 cells, the media was discarded, cells were scraped and collected. Cell pellets were obtained by centrifugation at 3000 rpm for 5 min at 4°C to remove the supernatants followed by one wash with cold 1xPBS. For total protein extraction, ice-cold lysis buffer K (20 mM phosphate buffer, 150 mM NaCl, 0.1% NP40, 5 mM EDTA, and proteinase inhibitors) was added to the washed cell pellets. Sonication was carried out at power level 4 for 5 sec, repeated 5 times, using a manual sonicator (model 100, Fisher Scientific) in a cold room on ice. This was followed by rotation at 4°C for 20 min. The solution was then centrifuged at 13,000 rpm for 10 min at 4°C and the resulting supernatants were transferred to a new tube. Protein concentration was determined using the Bradford method (Protein assay, 5000006, Bio-Rad).

For Western blotting, protein samples (30 µg) were separated on 8-12% SDS-PAGE gels, then transferred onto PVDF membranes (1620177, Bio-Rad). Subsequently, the membranes were blocked for 1 h at room temperature in TBS-T (0.1% Tween) containing 5% milk. The membranes were then incubated overnight at 4°C with primary antibodies, which were diluted in TBS-T containing 3% BSA and 0.3% sodium azide. After three washes in TBS-T, the membranes were incubated for 1 hour with anti-rabbit (NA934, GE Healthcare; RRID: AB_772206) antibody diluted in TBS-T containing 5% milk. After an additional three washes with TBS-T, protein bands were visualized using enhanced chemiluminescence (ECL) Clarity, ECL Clarity Max, or ECL Select Western blotting detection reagent (1705061 and 1705062, Bio-Rad, or RPN2235, GE Healthcare). The primary antibodies employed were: anti-ERRα (1:1000, ab76228, Abcam; RRID: AB_1523580), anti-HCFC1 (1:1000, A301-399A, Bethyl; RRID: AB_961012), and anti-β-actin (1:1000, ab8227, Abcam; RRID:AB_2305186). Immunoblots were analyzed using a ChemiDoc MP imaging system (Bio-Rad, Hercules, CA).

### 2.14 Mitochondrial respiration

The Oxygen Consumption Rates (OCRs) of Hepa 1-6 cells treated with either control siRNAs or siRNAs against ERRα and/or HCFC1 as described above were determined using an Agilent Seahorse XFe24 Extracellular Flux Analyzer (Agilent Technologies, Santa Clara, CA, USA) following the manufacturer’s guidelines. A day before the assay, cells were trypsinized and cell counts were adjusted to 30,000 cells per well in a XFe24 Cell Culture Miniplate. After allowing the cells to uniformly adhere to the well bottoms by incubating the plate at room temperature for 1 h, the plate was then transferred to a cell culture incubator (37°C with CO_2_) for overnight incubation. In parallel, the Seahorse sensor cartridge was hydrated by adding 1 ml of XF Calibrant to each well, followed by an overnight incubation at 37°C in an incubator without CO_2_. On the day of the assay (96 h post initial siRNA treatment), Seahorse assay media was preheated in a 37°C water bath for approximately 10-15 min. A total of 50 ml of Seahorse assay media was prepared using XF Base medium (103334-100, Agilent), with a final concentration of 25 mM glucose (G8769-100ML, Sigma), 1 mM sodium pyruvate (600-110-EL, Multicell), and 2 mM Glutamax (35050061, Thermo Fisher Scientific) with the pH adjusted to 7.4. The previous media was removed, and cells were washed once with 500 μl of assay media. Subsequently, assay media was added to all wells at a final volume of 500 µl. The plate was then placed in a 37°C incubator (without CO_2_) for 1 hour. Mitochondrial perturbators were loaded into ports A, B, and C of the Seahorse assay plate with 1 µM oligomycin (495455-10MG, Sigma), 80 µM 2,4-dinitrophenol (DNP, D198501; Sigma), and 0.5 µM rotenone (R8875-1G; Sigma) + 0.5 µM antimycin A (A8674-25MG; Sigma), respectively. Basal respiration was measured with 3 min of mixing and 3 min of measurement for three cycles. Measurements following subsequent injections of oligomycin, DNP, or rotenone + antimycin A were taken with 3 min of mixing, a 3-minute waiting period, and 3 min of measurement for a total of three measurement cycles. The values obtained were normalized to cell counts. Maximal OCRs were measured following the addition of the uncoupler DNP. Non-mitochondrial OCRs, residual respiration following addition of mitochondrial respiratory chain inhibitors rotenone and antimycin A, were subtracted from the basal and maximal OCRs for determination of mitochondrial respiration measurements. Results represent the average of four independent experiments.

### 2.15 Statistics and reproducibility

Unless otherwise specified, results are shown as means ± SEM with statistical analyses performed using GraphPad Prism 9 or 10. Significant differences were defined as *p < 0.05, **p < 0.01, ***p < 0.001, ****p < 0.0001 with the specific statistical tests detailed in the figure legends. The value of “n” for biological replicates represents either the number of individual mice used or independent biological replicates for cultured cell experiments.

### 2.16 Data availability

Mouse liver ERRα RIME MS proteomics data will be deposited to the ProteomeXchange Consortium via the PRIDE [38] partner upon publication. HCFC1 ChIP-seq data performed in Hepa 1-6 cells ± siRNA-mediated ERRα and HCFC1 knockdown as well as RNA-seq data from Hepa 1-6 cells ± siRNA-mediated knockdown of ERRα, HCFC1 or both will be deposited in NCBI’s Gene Expression Omnibus (GEO) upon publication. Publicly available ChIP-seq data analyzed in this study are available from GEO and summarized in Suppl. Table 4.

## 3. Results

### 3.1 ERRα RIME identification of associated proteins

In our initial investigation, we aimed to characterize the chromatin-bound interactome of ERRα in mouse liver by RIME. Given that the expression of ERRα is regulated in a circadian manner in the liver [39], RIME experiments were performed in triplicate at two distinct time points, ZT8 (day) and ZT24 (night) each with IgG controls, resulting in the comprehensive capture of 48 ERRα transcriptional partners through direct or indirect interactions (Figure 1A and Suppl. Table 1). The RIME experiments yielded a robust and consistent peptide coverage of ERRα across replicates and ZT times for a cumulative coverage of 60.7% (Figure 1B). A total of 40 ERRα interactors were identified at ZT24 as compared to 28 at ZT8, with 20 proteins commonly found at both time points examined (Figure 1C and Figure Supplemental 1A). The higher number of identified interactors at ZT24 was upheld by the generally increased normalized total precursor intensity (TPI) levels detected, highlighting the dynamic temporal nature of ERRα’s interactions and value of the experimental design to comprehensively map the ERRα hepatic interactome (Figure 1C). Upon comparison with established databases, 87.5% of identified interactors (42 of 48) were previously uncharacterized as proteins able to interact with ERRα (Figure 1D and Suppl. Table 2). Known ERRα coregulators including PROX1, PGC-1β, and NRIP1 were identified by RIME (Figure 1E), thus validating the effectiveness of the approach.

**Figure 1:**
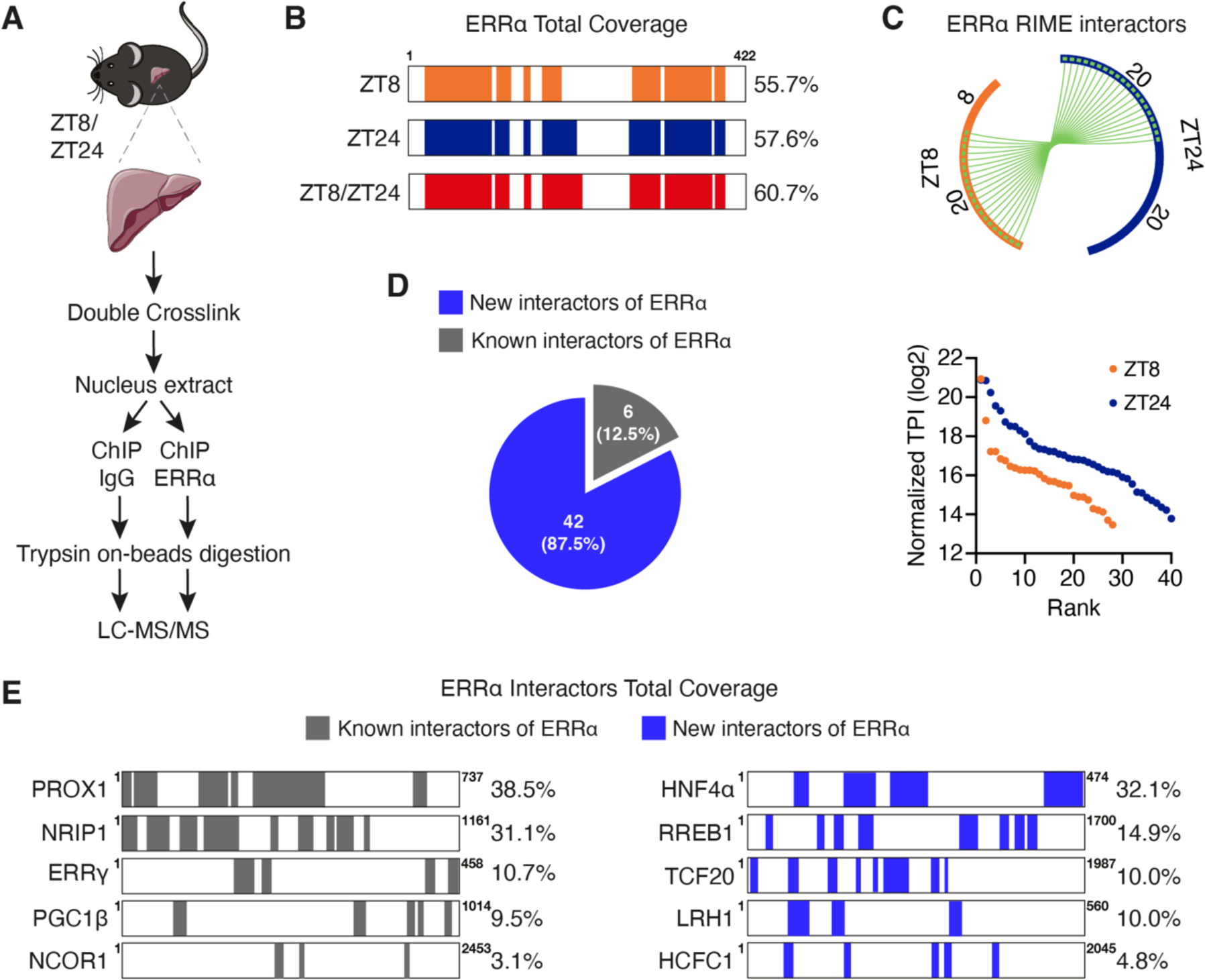
RIME identification of hepatic ERRα chromatin-bound interacting proteins. (A) Diagram illustrating the ERRα RIME protocol conducted in mouse livers isolated at ZT8 and ZT24. (B) RIME peptide coverage for ERRα at ZT8 (orange), ZT24 (blue) and the total coverage across both time points examined (red). (C) Top, Circos plot illustrating the relative number of ERRα-interacting proteins identified at the examined time points and their overlap. Orange arc, ZT8; blue arc, ZT24; connecting green lines, shared interactors at ZT8 and ZT24. Bottom, graph represents ranked normalized TPIs of identified hepatic ERRα RIME interactors at ZT8 and ZT24 using respecitve IgG controls (n=3). (D) Pie chart showing the proportion of identified RIME proteins previously known to interact with ERRα through examination of BioGRID, HuRI, IID, and STRING databases. (E) RIME peptide coverage for several of the ERRα identified interactors. Interactors in grey and blue denote known and newly identified partners, respectively. See also Suppl. Figure 1.

The total repertoire of 48 high confidence ERRα interactors included several transcription factors (e.g. CEBPA/B, RREB1) and nuclear receptors (e.g. HNF4α, NR5A2/LRH1) (Figure 2A). Functional interrogation by Metascape showed a strong enrichment of proteins involved in chromatin organization including multiple components of the Nucleosome Remodeling and Deacetylase (NURD) complex (MTA2, CHD4, GATA2A/B, and RBB4/7) also found tied to the top IPA enriched pathway associated with transcriptional repression (Figure 2B and 2C). Consistently, the major complex of functionally related ERRα interacting proteins was centered on the NURD complex known for its established role in deacetylation of histone tails and therefore its transcriptional repressive activity through chromatin condensation (Figure 2D). These findings underscore the repressive activity of ERRα in the liver, emphasizing its role in modulating gene expression through regulation of the chromatin state.

**Figure 2:**
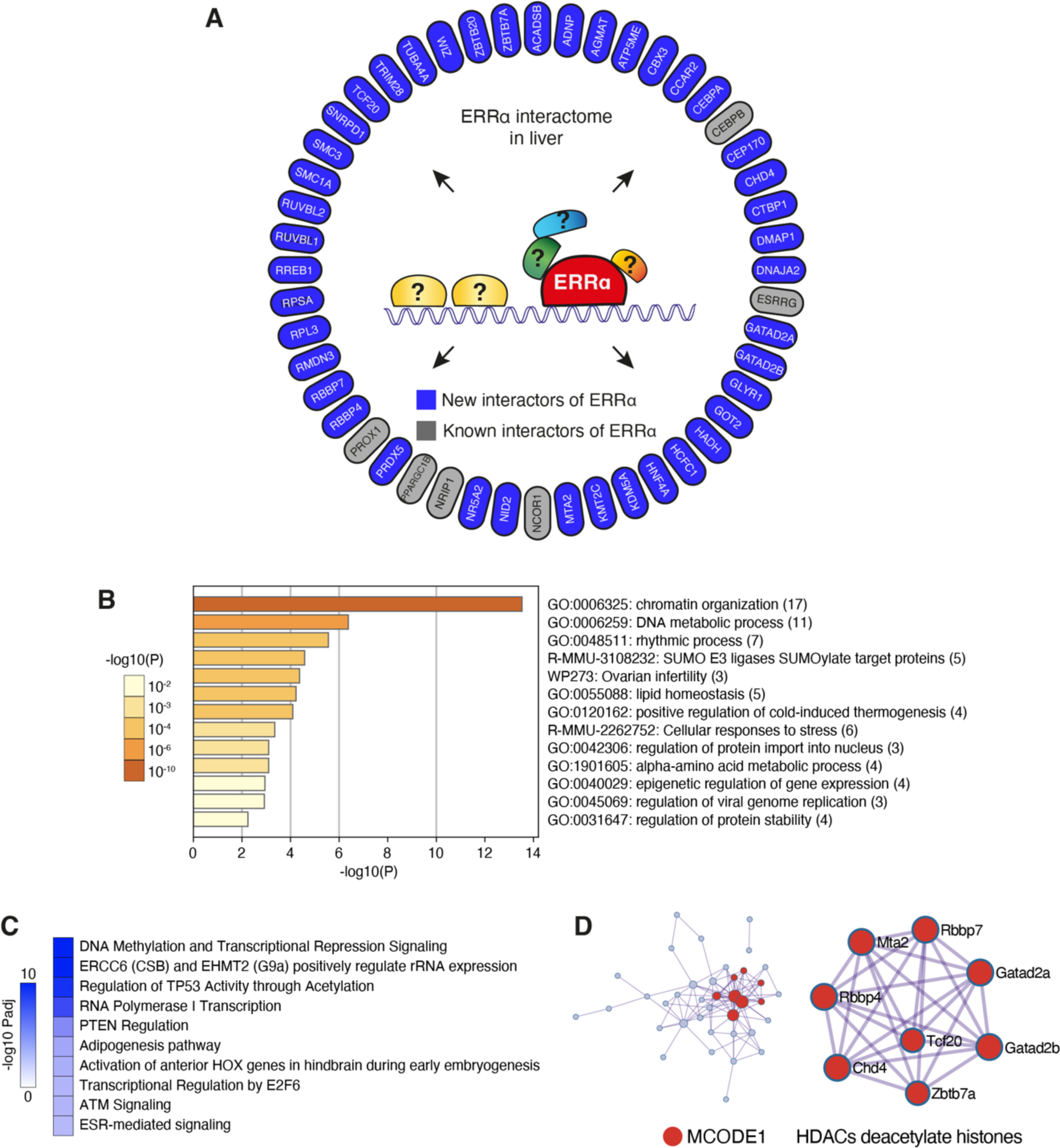
ERRα interactors are strongly associated with transcriptional repression. (A) Schematic displaying the total 48 high confidence proteomic interactors of ERRα identified by RIME in mouse liver. Proteins in gray and blue represent known and newly identified interactors, respectively. (B) Metascape enrichment analysis of the 48 ERRα interactors identified in (A) showing the top 13 statistically enriched biological pathway terms. (C) Top 10 significantly enriched IPA canonical pathways associated with the 48 ERRα interactors identified in (A). (D) Metascape protein-protein interaction network of the 48 ERRα interactors identified in (A) highlighting the enriched MCODE1 subcomplex associated with the term HDACs deacetylate histones in red.

### 3.2 Chromatin profiling analysis of ERRα and its identified cofactors

Following the discovery by RIME of 48 ERRα chromatin interactors including 42 that were previously unknown, we next examined the genomic crosstalk between ERRα and its transcriptional partners through integrative bioinformatics analyses. We initially assessed the availability of uniformly processed ChIP-seq profiles for the identified co-interactors within the ChIP-Atlas repository [40], determining that 50% (24 of 48) possessed at least one mouse-specific ChIP-seq dataset regardless of tissue or cellular context. Exploiting the functionality of the Enrichment Analysis tool within ChIP-Atlas, we next surveyed the overlap in binding profiles available for these 24 interactors with that of our previously generated mouse liver ERRα ChIP-seq dataset [34] also housed in ChIP-Atlas. Co-localization was observed with all 24 interactors (Figure 3A and Suppl. Table 4), strongly reinforcing the RIME findings. PROX1, found previously to act as an important corepressor of ERRα in liver [9; 39], was found to exhibit the greatest degree of overlap based on the average overlap from all available ChIP-seq datasets (Figure 3A). We next refined our investigation by focusing in on the 9 ERRα interactors with available liver/hepatocyte-specific ChIP-seq data and selected the best profiles for further analysis (Suppl. Table 4). Peak intersection analysis with the selected datasets by HOMER [32] showed generally increased overlaps with the mouse liver ERRα dataset (Figure 3B), reflected by the enrichment of cell-specific interactions and/or increased quality of the ChIP-seq datasets in this tissue/cellular context. Previously described interactors NCOR1, CEBPB, and PROX1 were found among the top colocalizing partners of ERRα (Figure 3B). Importantly, this computational analysis strongly supports the cooperation between ERRα and novel candidate interactors in liver: HNF4α, CEBPA, SMC3, TRIM28, HCFC1, and KMT2C (Figure 3B). Genome browser views of genes exhibiting ERRα colocalization with the subset of 9 interactors profiled in mouse liver/hepatocytes are shown in Figure 3C and Suppl. Figure 2.

**Figure 3:**
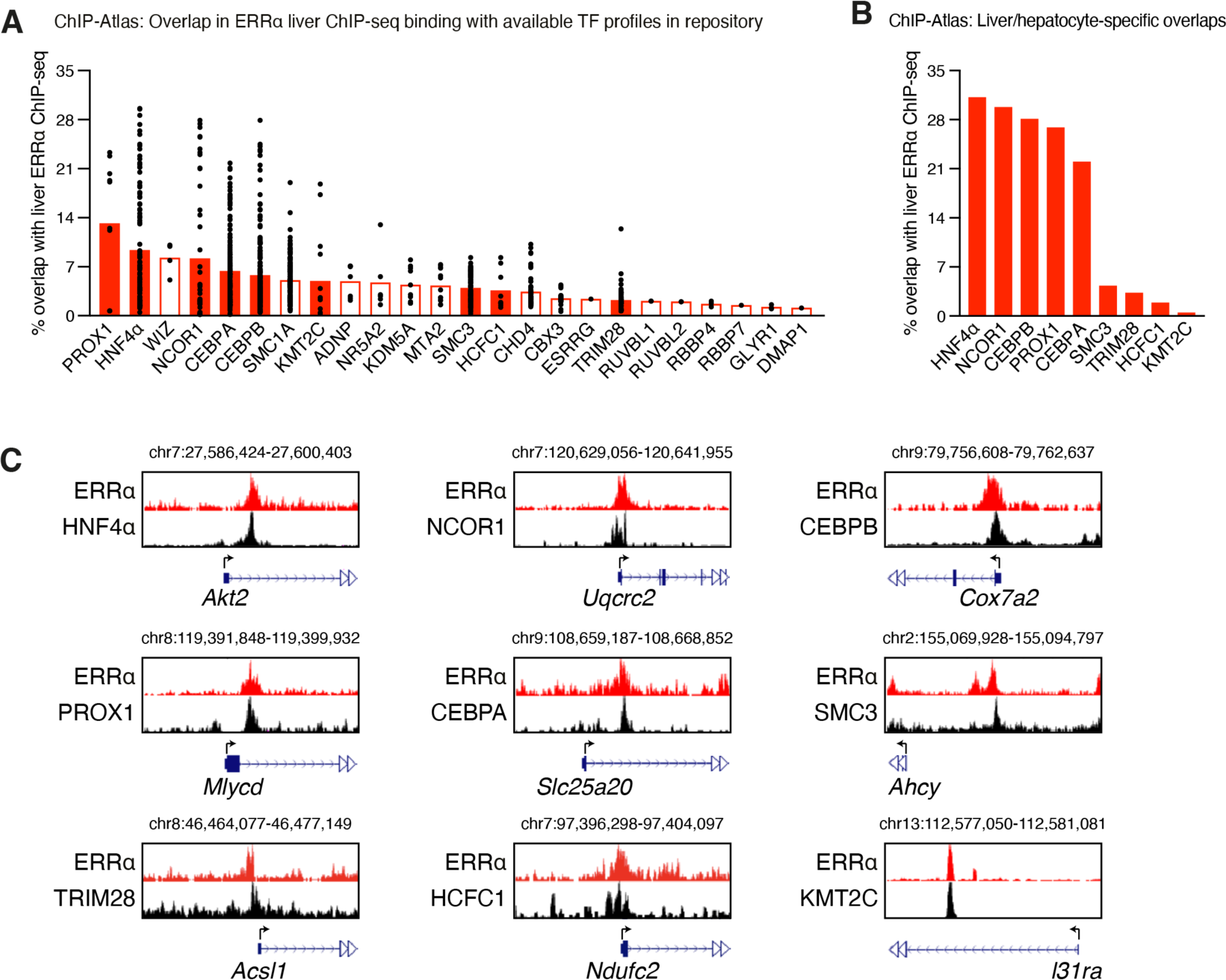
Genomic crosstalk between ERRα and its cofactors. (A) Graph illustrating the mean overlap between ERRα liver ChIP-seq profiles and the ChIP-seq profiles available for 24 of 48 RIME-identified ERRα interactors in the ChIP-Atlas repository across various mouse tissue/cellular contexts. Each black dot represents a ChIP-seq dataset. Of the 24 ERRα interactors with published ChIP-seq data, 9 had profiling conducted in mouse liver/hepatocytes as indicated by full red bars. (B) Graph showing the overlap between mouse ERRα liver ChIP-seq data and liver/hepatocyte-specific ChIP-seq profiles for 9 ERRα interactors identified in (A). (C) Examples of ChIP-seq tracks showing colocalization of ERRα and a subset of its RIME-identified interactors using profiles from (B). See also Suppl. Figure 2.

To define the functional relationship between ERRα and the refined 9-interactor subset in the liver, we next performed KEGG pathway enrichment analysis (p < 0.05) on the identified co-targeted genes with overlapping binding events found within ± 20kb of the TSSs (Figure 4A). Enriched pathways were found for all interrogated gene sets except for ERRα-KMT2C co-bound genes, likely due to the limited input size (Figure 4A).

**Figure 4:**
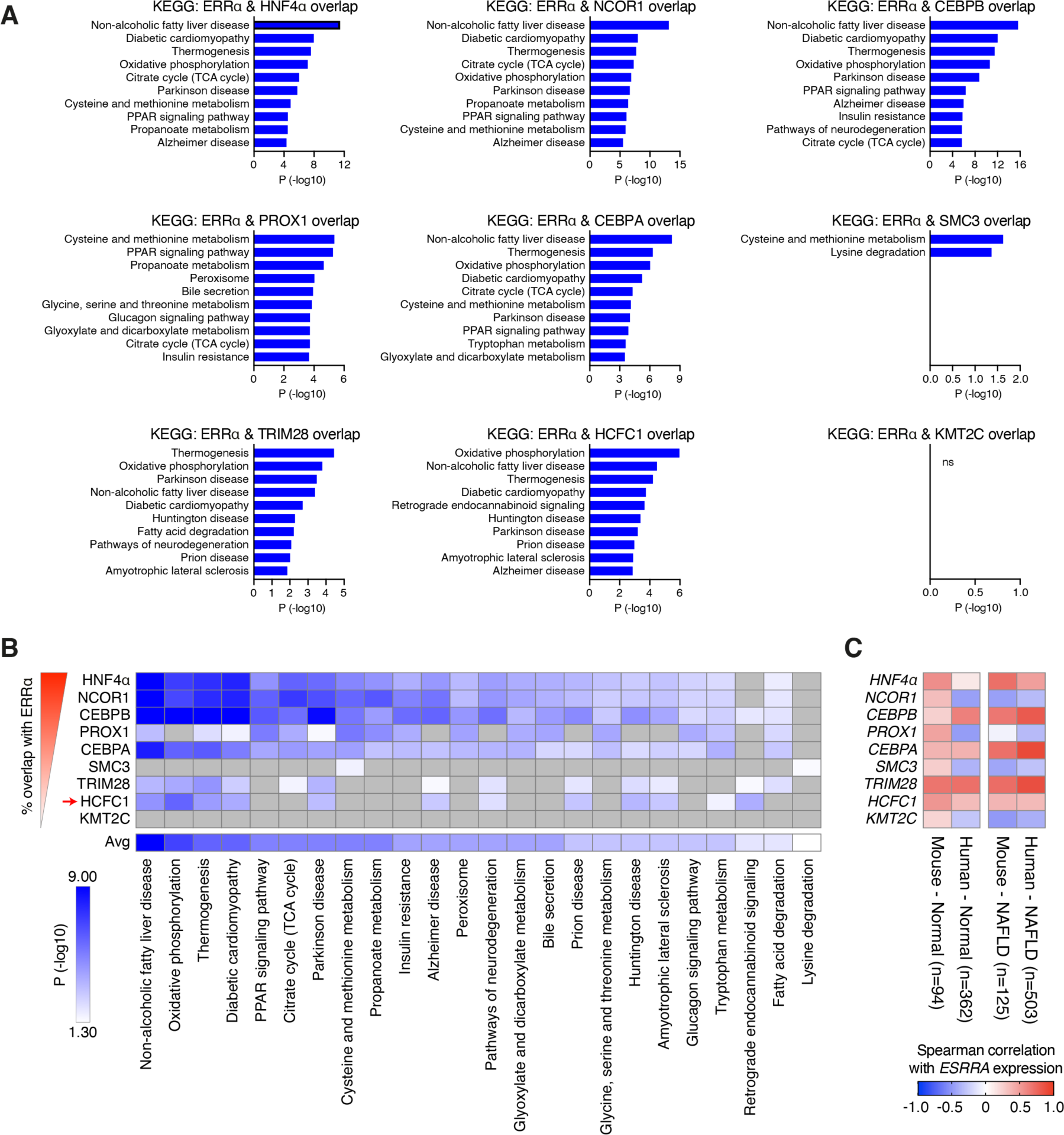
Functional relevance of genes co-bound between ERRα and its interactors. (A) Top 10 KEGG enriched pathways of genes with shared binding sites for ERRα and its 9 interactors with available liver/hepatocyte-specific profiles from Figure 3B. Genes were restricted to those with overlapping peaks found within ± 20kb of gene TSSs. (B) Heatmap recapitulating all top 10 KEGG pathways identified in (A) with pathways ranked by average significance across comparisons. Grey boxes signify that the pathway was not significantly enriched (p < 0.05) among that list of co-bound target genes. (C) Heatmap of Spearman correlation scores between *ESRRA* mRNA expression and the expression of 9 of its interactors in normal liver and NAFLD of both mouse and human origin.

Remarkably, further analysis of the top 10 KEGG pathways across comparisons revealed OXPHOS components as key co-occupied targets underscoring the top 4 most significant pathways based on mean significance, namely NAFLD (non-alcoholic fatty liver disease), OXPHOS, thermogenesis, and diabetic cardiomyopathy (Figure 4B). Interrogation of liver mouse and human expression profiles from a recently developed integrated liver expression atlas GepLiver [26] revealed similar correlation relationships between *ESRRA* expression and its 9 examined coregulators in mouse models and human patients with NAFLD (Figure 4C). Notably, negative correlations were found between *ESRRA* and its known transcriptional corepressors *NCOR1* and *PROX1*. Among the positive correlations, we noted the newly identified interactor *HCFC1*, found to correlate positively with *ESRRA* in the context of NAFLD as well as in the normal state (Figure 4C). HCFC1 is a transcriptional cofactor that modulates gene expression by interacting with various transcription factors [41]. Given that OXPHOS was the most enriched KEGG pathway among the ERRα-HCFC1 co-occupied genes in mouse liver (Figure 4B), a program at the center of a complex transcriptional network involving multiple coregulators of ERRα, we sought to detangle the transcriptional connection between ERRα and HCFC1. While ERRα and HCFC1 have been independently implicated in the transcriptional control of OXPHOS [42–44], no investigation has reported on their cooperation in the control of gene expression.

### 3.3 Validation of ERRα and HCFC1 interactions in mouse liver

To gain a deeper understanding of the relationship between ERRα and HCFC1 on chromatin, we executed an in-depth re-analysis of the published mouse liver ChIP-seq datasets from raw fastq files [34; 35]. Remarkably, despite targeting a common set of 3,684 genes within ± 20kb of gene TSSs, HOMER intersection of binding peaks revealed a small degree of overlap, identifying 601 co-occupied regions (Figure 5A and 5B). A significant proportion of these ERRα and HCFC1 overlapping peaks localized to promoter-TSS regions granted that 84% of the 601 common peaks were found ±1kb of TSSs (Figure 5C and 5D). The observed localization pattern of overlapping peaks was principally attributed to the known preferential occupancy of HCFC1 near TSSs, as found in our analysis in Figure 5C. Next, de novo motif analysis was performed on the three distinct peak sets identified in Figure 5B. As anticipated, the consensus ERR binding motif ERRE was found markedly enriched at ERRα-only sites (Figure 5E). Consistent with the fact that HCFC1 does not bind to DNA directly, enriched motifs for HCFC1-only peaks showed enrichment for transcription factors previously reported to interact with HCFC1, such as NRF1 [45] and THAP11 [41] (Figure 5E). Notably, the most enriched motif in ERRα and HCFC1 common peaks was the ERRE motif, suggesting a role for ERRα in recruiting HCFC1 to these sites (Figure 5E). Upon further inspection, ERRα and HCFC1 peak summits were found offset by a mean absolute distance of 364 bp with HCFC1 occupying regions closely upstream or downstream that of ERRα (Figure 5F and 5G). Despite this separation, we could confirm their specific colocalization using the same primer set by ChIP-qPCR assays in mouse liver at several target genes involved in OXPHOS (Figure 5H), their principal co-targeted pathway (Figure 4). Notably, ERRα and HCFC1 were found to co-occupy a significant proportion of all nuclear-encoded OXPHOS genes accounting for 38%, 75%, 56%, 50% and 78% of Complex I through V components, respectively (Figure 5H).

**Figure 5:**
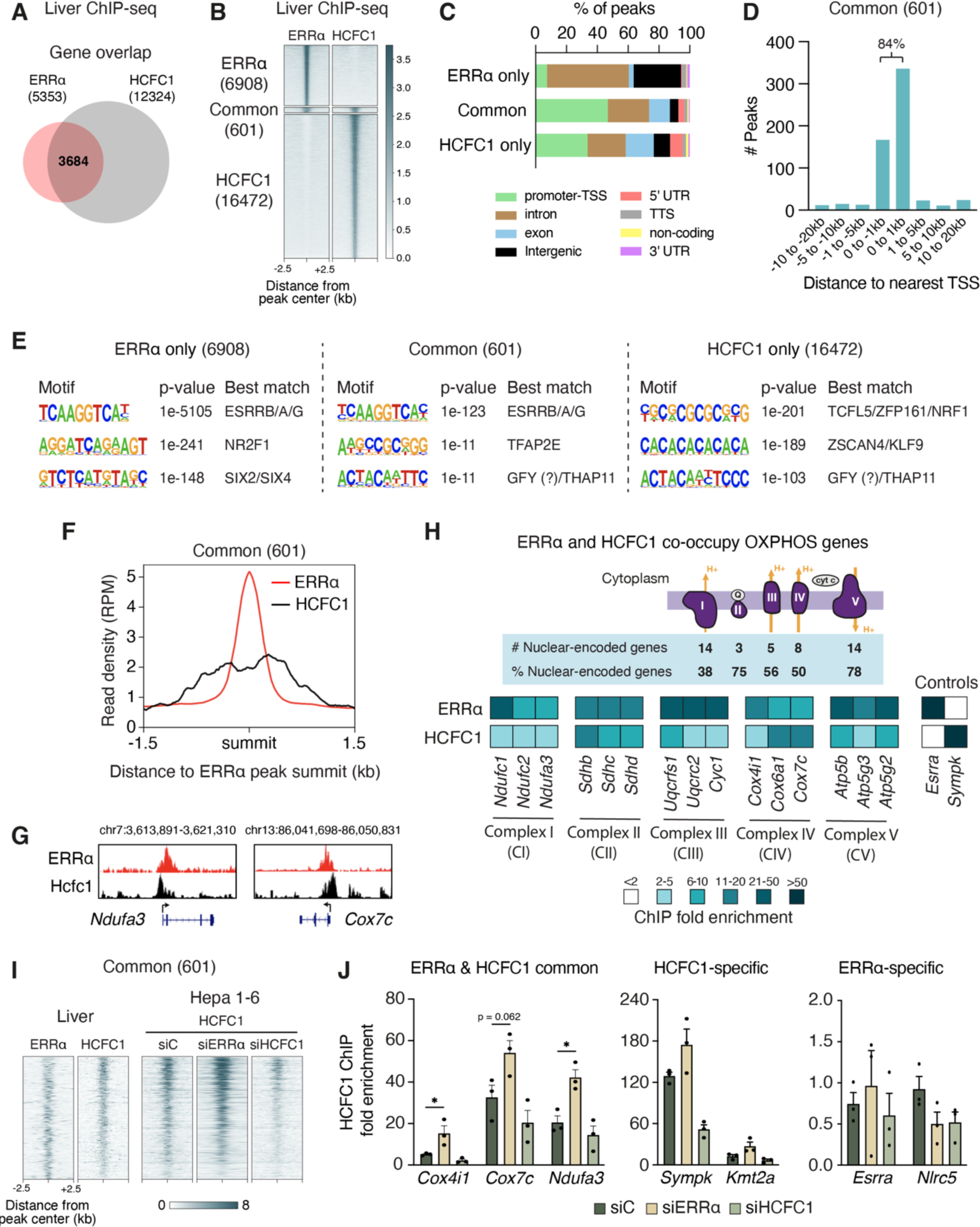
Liver ERRα and HCFC1 chromatin association. (A) Venn diagram illustrating the overlap in genes found targeted by ERRα (GSE43638) and HCFC1 (GSE115767) within ± 20kb of gene TSSs from ChIP-seq data in mouse liver. (B) Heatmap representing the binding signal intensities of resulting peak regions from ERRα (GSE43638) and HCFC1 (GSE115767) peak intersections based on peaks found ± 20kb of gene TSSs. (C) Genome annotations of peak locations for ERRα-specific, HCFC1-specific, and ERRα and HCFC1overlapping peaks (common) identified in (B) expressed as a percentage of the total peaks. (D) Distribution of peak positions for the 601 identified ERRα and HCFC1 overlapping peak regions in (B) relative to gene TSSs. (E) HOMER de novo motif analysis of peak lists in (B). (F) ChIP-seq read density graphs showing the distribution of HCFC1 peak summits relative to ERRα peak summits for the 601 shared regions identified in (B). (G) ChIP-seq tracks showing ERRα and HCFC1 colocalization at the indicated genes with the presence of an HCFC1 peak found directly upstream or downstream that of ERRα. (H) Top, schematic illustrating the co-occupancy of ERRα and HCFC1 at nuclear-encoded OXPHOS genes. The number of co-bound OXPHOS genes for each respiratory complex is shown and the proportion it represents among all nuclear-encoded genes. Bottom, heatmap showing mouse liver ChIP-qPCR enrichment values for ERRα and HCFC1 binding at OXPHOS co-targeted genes, validating their simultaneous occupancy at common sites. Control regions were used to validate ERRα- and HCFC1-uniquely bound regions. The data shown are from one experiment representative of two independent experiments. (I) Heatmap of ChIP-seq binding signal intensities for HCFC1 in Hepa 1-6 cells ± siRNA-mediated ERRα or HCFC1 knockdown at the 601 ERRα and HCFC1 co-occupied regions identified in mouse liver from (B). (J) HCFC1 ChIP-qPCR experiments in Hepa 1-6 cells ± siRNA-mediated ERRα or HCFC1 knockdown at established ERRα and HCFC1 co-bound sites as well as HCFC1- and ERRα-specific sites (n=3). Data (J) are presented as means ± SEM, *p < 0.05, unpaired two-tailed Student’s t test. See also Suppl. Figure 3.

Given the proximity in binding between ERRα and HCFC1 at 601 defined regions found enriched for the ERR binding motif ERRE, we next sought to establish whether HCFC1 recruitment to these regions is indeed ERRα-dependent. To this end, we performed HCFC1 ChIP-seq studies in the mouse liver cell line Hepa 1-6 ± siRNA-mediated ERRα knockdown (Figure 5I and Suppl. Figure 3A). Depletion of ERRα expression resulted in a visible augmentation in HCFC1 binding affinity at ERRα and HCFC1 co-targeted loci with a mean absolute fold change of 1.37 (Figure 5I). As a control, HCFC1 ChIP-seq conducted in cells with HCFC1 knockdown showed a general loss in binding, confirming the specificity of the antibody employed (Figure 5I). Surprisingly, however, while ChIP-qPCR experiments performed in triplicate confirmed a modest rise in HCFC1 recruitment to ERRα and HCFC1 co-targeted sites upon loss of ERRα (Figure 5J and Suppl. Figure 3B), there was also a trend for increased binding at HCFC1-specific sites (Figure 5J and Suppl. Figure 3C), suggesting that loss of ERRα expression results in the global enhancement of HCFC1 genomic binding. Indeed, quantification of HCFC1 protein levels showed a modest by significant 1.3-fold increase upon ERRα knockdown in Hepa 1-6 cells (Suppl. Figure 3D).

### 3.4 Loss-of-function studies reveal a differential dependency of ERRα on HCFC1 in the control of cellular respiration and proliferation

To further explore the transcriptional cooperation between ERRα and HCFC1, we next performed RNA-seq analyses in Hepa 1-6 cells subjected to siRNA-mediated ERRα and/or HCFC1 loss-of-function (Figure 6A and 6B). GSEA was employed for functional interrogation of transcriptome profiles to identify significantly deregulated gene signatures (FDR < 0.25, n =3). While knockdown of ERRα predominantly resulted in a downregulation of Hallmark gene sets, loss of HCFC1 showed a more mixed regulatory modulation of signatures (Figure 6C). HCFC1 ablation discernably dominated the phenotype of cells with combined ERRα and HCFC1 deficiency (Figure 6C), reflecting the importance of HCFC1 as a transcriptional cofactor. Among the gene signatures found commonly and independently regulated by loss of ERRα or HCFC1 alone, we noted OXPHOS and the cell cycle-related term G2M_CHECKPOINT (Figure 6D). ERRα ablation significantly upregulated the OXPHOS gene signature as opposed to HCFC1 deficiency which showed a significant downregulation of the gene set with loss of both factors nulling the effects of each knockdown alone (Figure 6D and 6E). Several of the differentially regulated OXPHOS genes were previously validated for ERRα and HCFC1 co-localization namely *Atp5g3*, *Cox4i1*, *Cox7c*, and *Ndufa3* (Figure 5H). In contrast to the observed differential regulation of ERRα and HCFC1 on mitochondrial OXPHOS genes, both factors displayed a positive transcriptional action on cell cycle genes with their loss-of-function leading to their significant downregulation (Figure 6D and 6F). We next explored whether HCFC1 loss functionally impacts ERRα regulation of OXPHOS and cell cycle control by measuring the rates of Hepa 1-6 mitochondrial oxygen consumption and cellular growth. Importantly, the observed upregulation of basal and maximal rates of mitochondrial respiration in ERRα-deficient cells were abolished in cells with concomitant loss of HCFC1 (Figure 6G). On the other hand, loss of HCFC1 further reduced the repressive effect of ERRα knockdown on cell proliferation (Figure 6H). These biological phenotypes largely mirrored the dysregulation of OXPHOS, and cell cycle gene signatures observed upon ERRα and/or HCFC1 knockdown (Figure 6C-6F) and highlight the specificity of HCFC1 as an influencer of ERRα transcriptional control.

**Figure 6:**
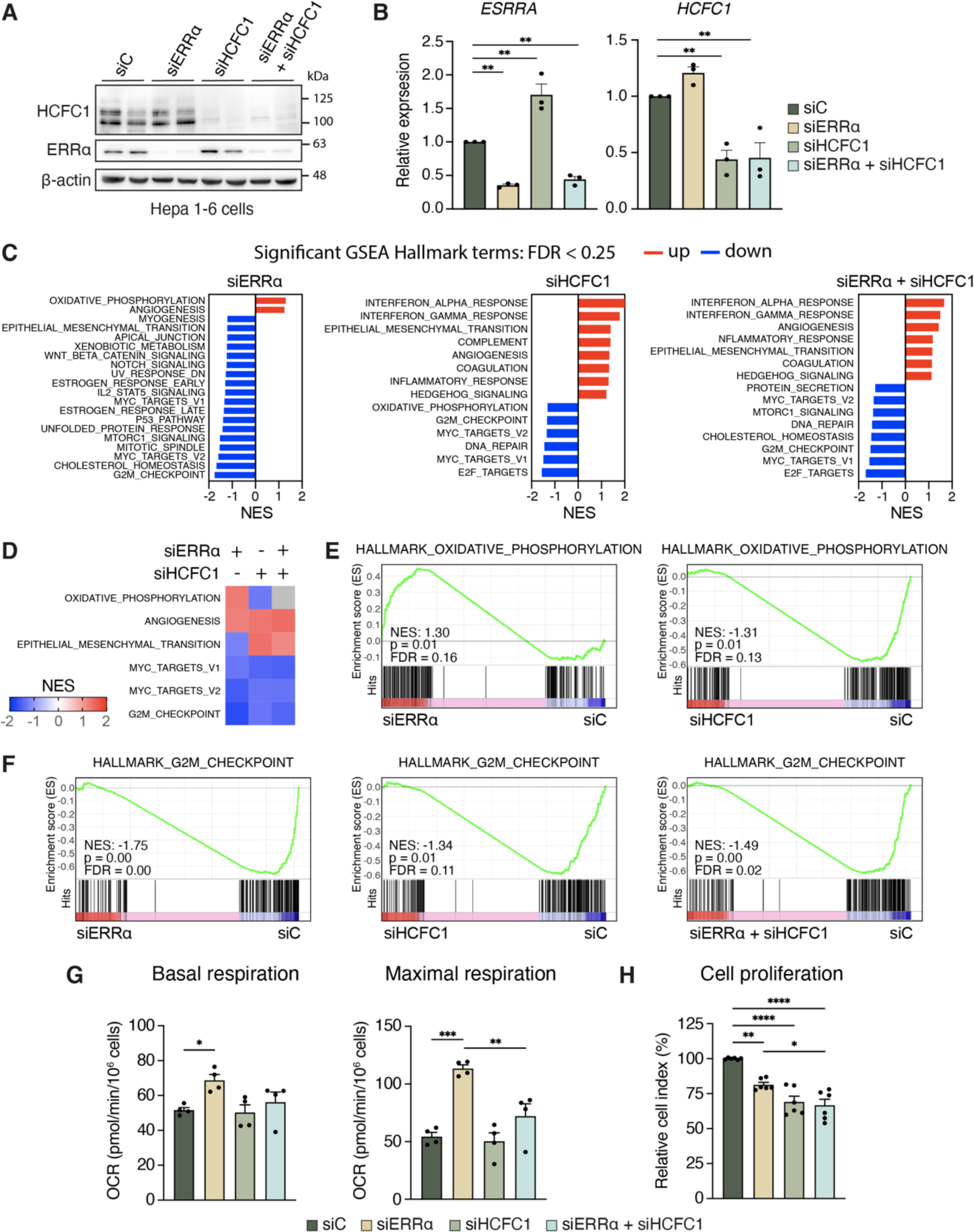
ERRα transcriptional regulation of OXPHOS genes is HCFC1-dependent. (A,B) Evaluation of the effectiveness of a pool of siRNAs against ERRα and/or HCFC1 in Hepa 1-6 cells at the protein (n=2) (A) and mRNA (n=3) (B) level by Western blot and RT-qPCR analyses, respectively. (C) GSEA analysis of Hepa 1-6 transcriptomes upon siRNA-mediated ERRα and/or HCFC1 knockdown (n=3). Significantly enriched gene sets with an FDR < 0.25 are shown. Bars represent the normalized enrichment score (NES): red and blue bars signify that the gene set is upregulated and downregulated upon the indicated knockdown, respectively. (D) Heatmap summarizing the common GSEA hallmark terms found independently modulated by the loss of ERRα or HCFC1 alone in (C). (E,F) GSEA enrichment plots for the significantly enriched hallmark term OXPHOS (E) and cell cycle-related term G2M_CHECKPOINT (F) shown in (D). (G) Basal and maximal. mitochondrial respiration rates of Hepa 1-6 cells treated with siRNAs against ERRα and/or HCFC1 (n=4) measured using a Seahorse Flux Analyzer. (H) Assessment of cell proliferation after a 72 h siRNA-mediated knockdown of ERRα and/or HCFC1 in Hepa 1-6 cells (n=6) expressed as a percentage relative to the control knockdown (siC). Data (B, G, and H) are presented as means ± SEM, *p < 0.05, **p < 0.01, ***p < 0.001, ****p < 0.0001, one-way ANOVA with Tukey’s post-hoc analysis.

## 4. Discussion

In this study, the application of RIME has yielded a comprehensive elucidation of the ERRα chromatin-bound interactome, a first for an orphan nuclear receptor, revealing 48 proteins that were detected in mouse liver through complex assembly or neighboring chromatin contacts. PGC-1β, PROX1, NCoR1, NRIP1, ERRγ, and CEBPB were previously recognized as ERRα cofactors [9; 10; 21; 46-48], reinforcing the validity of our findings. The discovery of 42 novel ERRα interacting proteins including transcription factors, cofactors and modifiers of chromatin structure expands our current understanding of its molecular network, substantiating the significance of the findings.

First, we performed RIME analyses on livers harvested at ZT24 and ZT8, shown in our previous report to concord with the highest and lowest diurnal protein levels of ERRα, respectively [39]. These findings alluded to a possible temporal variation in the ERRα interactome or at the least that its increased abundance at ZT24 would facilitate the enrichment of ERRα and thereby detection of its interactors by MS. Indeed, our analyses revealed a more extensive interactome of ERRα at ZT24 compared to ZT8, underscored by a general increase in binding affinities, despite achieving a similar coverage in ERRα protein, warranting future investigation into the temporal dynamics of ERRα interactions with these proteins and their implications on a mechanistic level.

Second, a significant observation from our study was the enrichment of identified ERRα interactors associated with repressive actions and lack thereof of coactivators. Intriguingly, while PGC-1α is regarded as the main coactivator of ERRα, only PGC-1β was found in our RIME study. Examination of gene expression profiles in GepLiver [26] revealed a 1.8-fold higher median expression level of the PGC-1β vs PGC-1α encoding genes in the normal mouse liver. This is in stark contrast to the nearly 11-fold higher PGC-1α vs PGC-1β median levels observed in the normal human liver. Although ERRα is commonly associated with transcriptional activation and the induction of target genes, its role within the liver is context-dependent, displaying both activating and repressive effects on gene expression [3]. This repression is facilitated through various mechanisms, including competition with other nuclear receptors or transcription factors, interactions with corepressors and the recruitment of chromatin-modifying complexes [16; 49]. For example, we have shown recently that ERRα and PPARα compete for shared sites in the context of fatty liver and NAFLD [8]. While PPARα was not identified in our RIME experiments in the normal mouse liver, it prompts interest for the elucidation of the ERRα interactome under pathological conditions for comparison to its associations in the normal state. Our reported interactions found by RIME with corepressors such as PROX1, NCoR1 and NRIP1, highlight an important function for ERRα in hepatic transcriptional repression [9; 21]. The discovery of multiple components of the NuRD complex known to play a central role in gene repression through chromatin remodeling and histone deacetylation, suggests that the NuRD compressor complex likely participates in shaping the repressive function of ERRα.

Third, our computational analyses revealed that ERRα transcriptional regulation of OXPHOS, a main target of ERRα metabolic function, involves a complex transcriptional network encompassing multiple coregulators including HCFC1 whose expression was found to correlate positively with that of ERRα in liver. By acting as a transcriptional coregulator, HCFC1 has been shown to play a crucial role in regulating the expression of genes involved in physiological processes such as cell growth, proliferation, and differentiation [50–52] and notably was implicated in the regulation of the insulin receptor binding to DNA [53]. While ERRα and HCFC1 have been independently implicated in the transcriptional control of metabolic processes such as OXPHOS [42–44], lipogenesis [54; 55], and gluconeogenesis [43; 56], cooperative interactions in metabolic control had never been reported. Through functional genomics analyses, we demonstrated that HCFC1 acts as a cofactor for ERRα in the transcriptional regulation of OXPHOS gene expression, revealing a new regulatory axis in the control of cellular energy metabolism. Together, our findings thus contribute valuable insights into the multifaceted roles and regulatory mechanisms orchestrated by ERRα in the context of liver biology and emphasize the importance of delving into the molecular intricacies of these interactions within the broader framework of liver function, homeostasis, and the disease state.

## 5. Conclusions

In conclusion, our investigation reveals a comprehensive profile of the ERRα interactome within the normal liver by introducing 42 previously unidentified proteins as key actors in an intricate interconnected transcriptional network. Within this complex interplay, we show specificity in the genomic associations between ERRα and several of its interactors and further demonstrate that HCFC1 cooperates with ERRα in the control of OXPHOS gene expression.

## Author contributions

Conception and design: C.S. and V.G.; Development of methodology: C.S.; Acquisition of data: C.S., E.P., P.H., S.K., R.B., and C.G.; Analysis and interpretation of data: C.S., C.R.D., and V.G.; Writing of the manuscript: C.S., C.R.D., and V.G.; Study supervision: V.G.

## Acknowledgements

We thank all members of the V.G. laboratory for helpful discussions and the staff of the proteomic facility at the Institut de Recherches Cliniques de Montréal, especially Denis Faubert and Josée Champagne. We thank the Bioinformatics Core Technology Platform, especially Alain Pacis. This study was supported by a Terry Fox Research Institute Team Grant (PPG-1091) and a Foundation Grant from the Canadian Institutes of Health Research to V.G. (FDT-156254). C.S was supported by a Canderel Scholarship.

## APPENDIX A

SUPPLEMENTARY DATA.

## Abbreviations

ChIP: chromatin immunoprecipitation
ChIP-seq: chromatin immunoprecipitation sequencing
DEGs: differentially expressed genes
ERR: estrogen-related receptor
ERRE: ERR response element
HCFC1: host cell factor C1
NAFLD: non-alcoholic fatty liver disease
NURD: nucleosome remodeling and deacetylase
OXPHOS: oxidative phosphorylation
OCR: oxygen consumption rate
PCR: polymerase chain reaction
PGC-1: peroxisome proliferator-activated receptor γ coactivator 1
PROX1: prospero homeobox protein 1
RIME: rapid immunoprecipitation mass spectrometry of endogenous proteins
RIP140: receptor interactor protein 140
TPI: total precursor intensity
TSS: transcriptional start site
WT: wild-type
ZT: zeitgeber time

