## Supplemental Info for "Identification of a chromatin-bound ERRα interactome network in mouse liver"

**Supplemental Figures 1-3**

**Supplemental Tables 1-7 (Excel Files)**

### Supplemental Figure 1

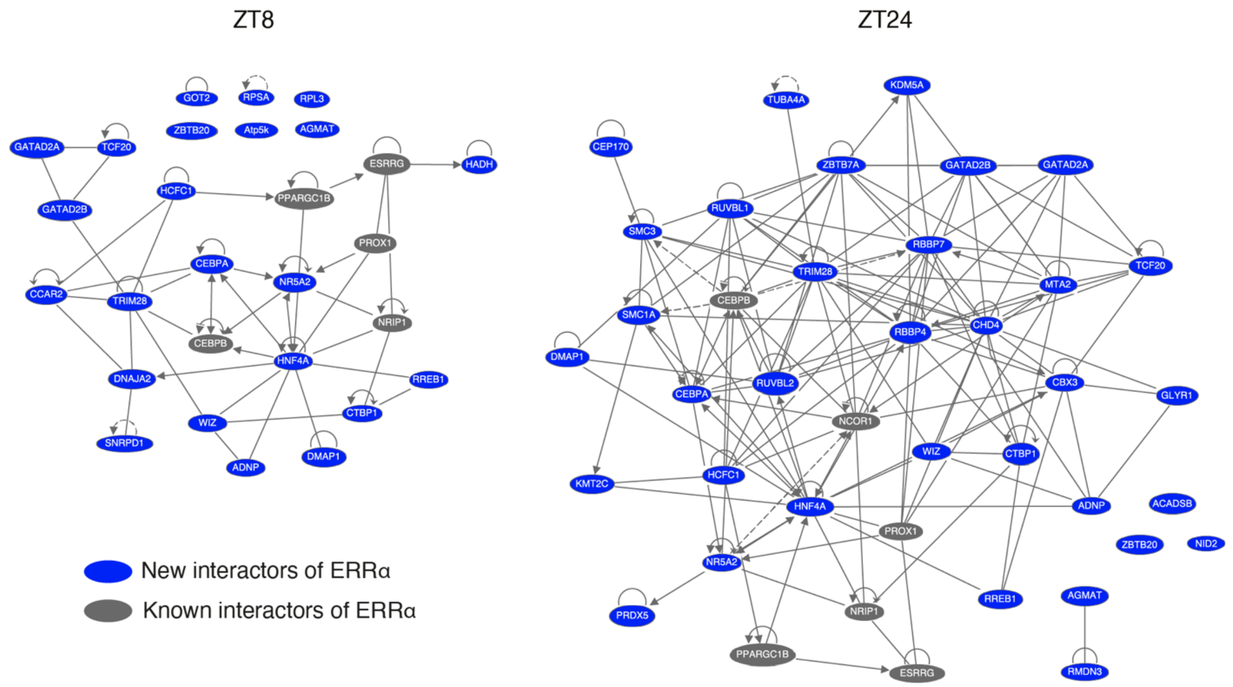


**Supplemental Figure 1:** IPA protein-protein interaction networks of the identified ERRα RIME interactors at ZT8 and ZT24. Connections represent either direct or indirect relationships from IPA's knowledgebase. See also Figure 1.

### Supplemental Figure 2

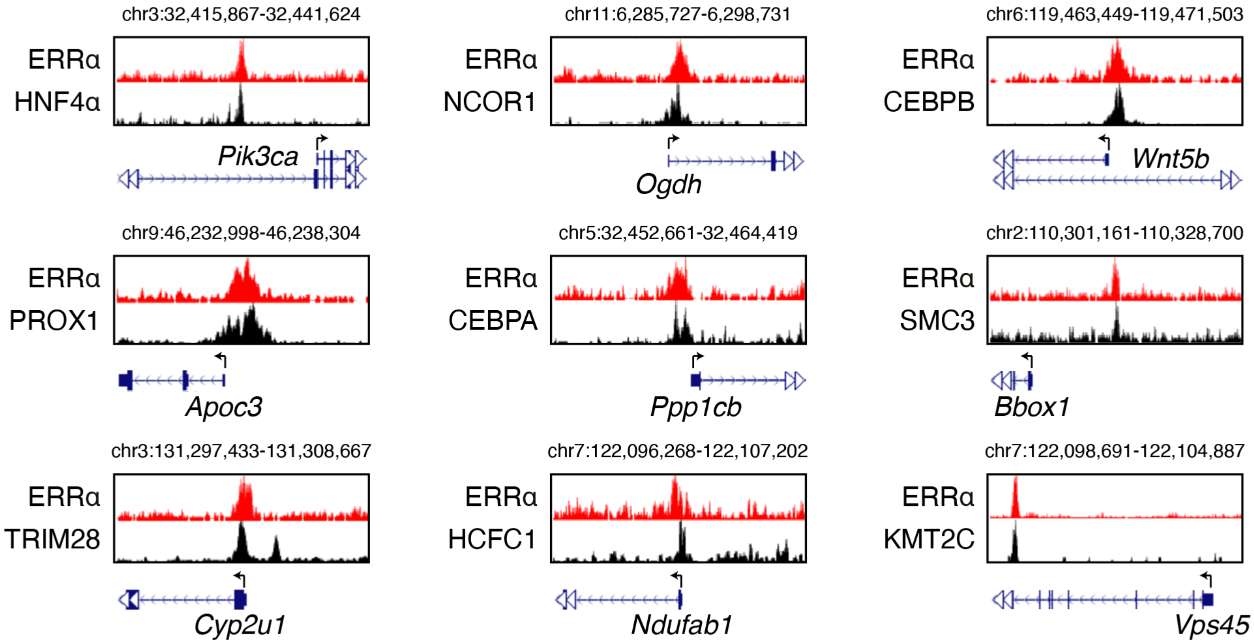


**Supplemental Figure 2:** Examples of ChIP-seq tracks showing colocalization of ERRα and a subset of its RIME-identified interactors using liver/hepatocyte-specific profiles from Figure 3B. See also Figure 3.

### Supplemental Figure 3

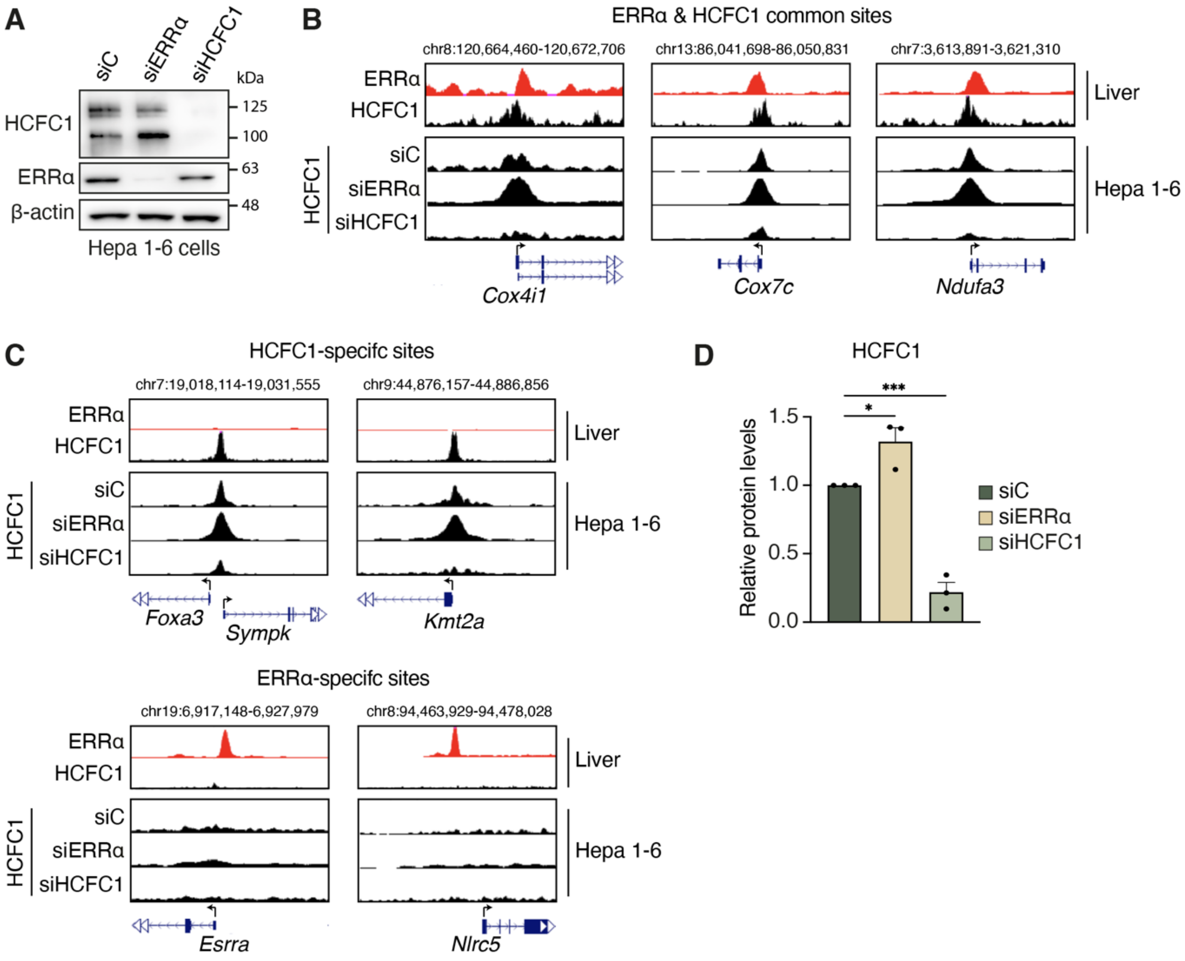


**Supplemental Figure 3:** (A) Representative Western blot of Hepa 1-6 cells ± siRNA-mediated ERRα or HCFC1 knockdown. (B,C) ChIP-seq tracks showing HCFC1 binding in Hepa 1-6 cells ± siRNA-mediated ERRα or HCFC1 knockdown at established ERRα and HCFC1 co-targeted OXPHOS genes (B) as well as at HCFC1- and ERRα-specific regions (C) identified from re-analysis of mouse liver ERRα (GSE43638) and HCFC1 (GSE115767) ChIP-seq data. (D) Quantification of HCFC1 protein levels in Hepa 1-6 cells ± siRNA-mediated ERRα or HCFC1 knockdown from triplicate independent experiments. Data (D) are presented as means ± SEM, *$p < 0.05$, ***$p < 0.001$, one-way ANOVA with Tukey's post-hoc analysis. See also Figure 5.

**Supplemental Table 1:** Identified ERRα chromatin-interacting proteins by RIME analysis in mouse liver at ZT8 and ZT24.

**Supplemental Table 2:** Known ERRα-interacting proteins compiled from BioGRID, HuRI, IID, and STRING databases.

**Supplemental Table 3:** Mouse primers used in this study.

**Supplemental Table 4:** Overlap of mouse liver ERRα ChIP-seq with available profiles of a subset of ERRα RIME interactors by ChIP-Atlas.

**Supplemental Table 5:** List of 601 genomic regions with overlapping ERRα and HCFC1 ChIP-seq peaks within ± 20kb of gene TSSs identified in mouse liver.

**Supplemental Table 6:** HCFC1 ChIP-seq in Hepa 1-6 cells ± ERRα or HCFC1 siRNA-mediated knockdown for 96 h.

**Supplemental Table 7:** RNA-seq analysis of Hepa 1-6 cells ± ERRα and/or HCFC1 siRNA-mediated knockdown for 96 h.
